# Evidence for white matter intrinsic connectivity networks at rest and during a task: a large-scale study and templates

**DOI:** 10.1101/2024.12.16.628798

**Authors:** Vaibhavi Itkyal, Armin Iraji, Kyle M. Jensen, Theodore J. LaGrow, Marlena Duda, Jessica A. Turner, Jingyu Liu, Lei Wu, Yuhui Du, Jill Fries, Zening Fu, Peter Kochunov, A Belger, J M Ford, D H Mathalon, G D Pearlson, S G Potkin, A Preda, T G M van Erp, K Yang, A Sawa, K Hutchison, E A Osuch, Jean Theberge, C Abbott, B A Mueller, Jiayu Chen, J Sui, Tulay Adali, Vince D. Calhoun

## Abstract

Understanding white matter (WM) functional connectivity is crucial for unraveling brain function and dysfunction. In this study, we present a novel WM intrinsic connectivity network (ICN) template derived from over 100,000 fMRI scans, identifying 97 robust WM ICNs using spatially constrained independent component analysis (scICA). This WM template, combined with a previously identified gray matter (GM) ICN template from the same dataset, was applied to analyze a resting-state fMRI (rs-fMRI) dataset from the Bipolar-Schizophrenia Network on Intermediate Phenotypes 2 (BSNIP2; 590 subjects) and a task-based fMRI dataset from the MIND Clinical Imaging Consortium (MCIC; 75 subjects). Our analysis highlights distinct spatial maps for WM and GM ICNs, with WM ICNs showing higher frequency profiles. Modular structure within WM ICNs and interactions between WM and GM modules were identified. Task-based fMRI revealed event-related BOLD signals in WM ICNs, particularly within the corticospinal tract, lateralized to finger movement. Notable differences in static functional network connectivity (sFNC) matrices were observed between controls (HC) and schizophrenia (SZ) subjects in both WM and GM networks. This open-source WM NeuroMark template and automated pipeline offer a powerful tool for advancing WM connectivity research across diverse datasets.

## Introduction

Functional magnetic resonance imaging (fMRI) has enabled us to observe the functioning of the human brain non-invasively (Nozais et al., 2023; Ogawa et al., 1990). By exploiting the blood-oxygenation-level-dependent (BOLD) effect, fMRI captures changes in blood flow related to neural activity, providing insights into brain function under both task-based and resting-state conditions. Task-based fMRI has been instrumental in mapping brain regions involved in specific cognitive and sensory processes by comparing brain activity during task performance to baseline states. Resting-state fMRI (rsfMRI) offers a complementary perspective by examining the brain’s activity in the absence of overt tasks (Biswal et al., 1995; M. H. Lee et al., 2013). This approach reveals spontaneous low-frequency oscillations in the BOLD signal, which reflect intrinsic functional connectivity networks (ICNs) and has significantly advanced our understanding of the brain’s functional organization.

Traditionally, the focus of fMRI studies has been predominantly on gray matter (GM), which is known to exhibit robust and well-characterized BOLD signals. The examination of white matter (WM) functional connectivity, however, has been relatively underexplored. In particular, WM has received less focus in functional studies due to its lower blood flow and volume compared to GM (Gore et al., 2019; Helenius et al., 2003; Lv et al., 2018; Rostrup et al., 2000) and the biological mechanisms underlying the BOLD signal in white matter remains under investigation (Logothetis & Wandell, 2004; Schilling et al., 2023). However, recent studies demonstrate that BOLD signals in WM are detectable and exhibit characteristics similar to those in GM, albeit weaker and with longer latencies (Gawryluk et al., 2014; Gore et al., 2019; Schilling et al., 2023). Emerging evidence suggests that BOLD signals in WM are not only detectable but also modulated by neural activity in interconnected GM regions (Gore et al., 2019). Despite the lower blood flow observed in WM compared to GM, the oxygen extraction fractions in WM are similar to those in GM (Raichle & Snyder, 2007). For instance, WM BOLD responses to stimuli are measurable and WM regions can show reliable activations during functional tasks (Gore et al., 2019). Moreover, the higher glia-to-neuron ratio in WM (Hofmann et al., 2017) implies a significant metabolic activity that supports the maintenance of myelin and neural communication, further supporting the notion that WM plays an active role in brain function (Gore et al., 2019). These findings highlight the importance of re-evaluating WM’s role in functional connectivity studies (Forkel et al., 2022) and suggest that current analytical methods may not fully capture the contributions of WM (Hu et al., 2023; Wang et al., 2022, 2023).

To address these gaps, our study leverages advanced techniques to explore both resting-state and task-based fMRI data, providing a more comprehensive understanding of WM functional connectivity. We introduce a novel WM ICN template derived from an extensive dataset of over 100,000 fMRI scans, building on methodologies previously utilized in (Iraji et al., 2023) as they ensure replicability and generalizability across diverse populations, enhancing their clinical utility. These templates were derived using multi-model-order independent component analysis (ICA) (Iraji et al., 2022, 2023), and then used as spatial priors within a spatially constrained ICA (scICA) pipeline called NeuroMark (Du et al., 2020; Iraji et al., 2023). ICA has been validated extensively in cortical studies, proving effectiveness in identifying inter-regional relationships and functional connectivity (Dini et al., 2024; Franco et al., 2013; Hassanzadeh et al., 2024; Xiong et al., 2020). By applying scICA to the whole brain data and using WM and GM templates, we can extend these capabilities to explore the functional connectivity patterns within WM and GM, thereby enhancing our understanding of their roles in brain function. Additionally, ICA can identify functionally related structures within WM that may correspond to specific tracts identified from diffusion magnetic resonance imaging (MRI), providing insights into structural-functional relationships (Tang et al., 2017). The NeuroMark ICA pipeline enhances these capabilities by offering a robust and adaptable framework for analyzing functional connectivity (Du et al., 2020). Unlike atlas-based methods that rely on fixed regions of interest (ROIs), NeuroMark’s data-driven approach adapts to individual scans, capturing subject-specific information while maintaining inter-subject correspondance. This flexibility enhances the reproducibility and comparability of studies, providing reliable imaging markers across different subjects and datasets. NeuroMark’s ability to integrate structural and functional data further extends its utility, allowing for a comprehensive analysis of brain function and structure.

Our study first identifies a robust set of WM ICNs to build a new WM template using a multiscale ICA approach over 100,000 fMRI scans (Iraji et al., 2023) (see Methods). This novel WM ICN template provides reliable priors for extracting WM ICNs from individual scan. Next we combine this template with the GM template (Iraji et al., 2023) and apply them to analyze resting-state and task-based fMRI data to investigate their spatial maps and functional connectivity of WM and GM separately as well as combined GM and WM. The resting-state data from the Bipolar-Schizophrenia Network on Intermediate Phenotypes 2 (BSNIP2) dataset (Tamminga et al., 2014), comprising 590 subjects, allows for a detailed exploration of spatial activation, functional network connectivity (FNC), and time spectral properties of the templates. We examined the FNC matrices for both WM and GM. We found significant differences in WM FNC between controls and schizophrenia patients. In addition, we applied these templates through the NeuroMark pipeline to task-based fMRI data during a sensorimotor task from the MIND Clinical Imaging Consortium (MCIC) dataset (Gollub et al., 2013), with 75 subjects. We show that WM ICNs manifest stimulus-evoked BOLD signals within the corticospinal tract during a sensorimotor task. The WM task-effects also show increased latency relative to the GM task-effects. This work can help us further understand functional connectivity in WM, integrating both intrinsic resting-state fluctuations and task-evoked activations. The analysis reveals distinct anatomical and functional differences between GM and WM ICN templates, with WM ICNs exhibiting higher-frequency peaks around 0.06 Hz. Modularization of the sFNC matrices highlights 13 WM and 14 GM domains, with notable inter-domain connectivity patterns, especially in the frontal, insular, and temporoparietal regions. In schizophrenia, significant reductions in WM connectivity, particularly in the insular and subcortical regions, are observed, while GM connectivity shows both hyperconnectivity and reductions, possibly reflecting compensatory mechanisms. These schizophrenia-related alterations underscore the importance of studying BOLD-related WM connectivity. To facilitate further research, we release the multi-model order ICA template and fully automated NeuroMark ICA pipeline for further study of this important topic.

## Methods

### Overall analysis framework

Our study employed advanced neuroimaging and analytical techniques to create a WM ICN template and further investigate functional connectivity patterns in SZ and controls (HC) using both resting-state and task-based fMRI data. We generated a WM template consisting of 97 robust and replicable WM ICNs using multiscale independent component analysis (Iraji et al., 2023), which were further combined with a 105 GM ICN template (Iraji et al., 2023). Multi-objective optimization ICA with reference (MOO-ICAR, (Du et al., 2020)) was used to implement scICA from the GIFT toolbox (https://trendscenter.org/software/gift/) (V. D. Calhoun et al., 2001; Iraji et al., 2021), with WM and GM templates as the reference. We calculated subject-specific ICNs spatial maps (SM) and time courses (TC) and computed static functional network connectivity (sFNC) measures for both GM and WM ICNs (summarized below). We also transformed time series data into the frequency domain to obtain power spectra and modularized connectivity matrices to identify functional modules. Task-based analysis focused on identifying neural correlates of cognitive tasks through scICA and temporal sorting. Our comprehensive approach, integrating GM and WM connectivity analyses, provides a nuanced understanding of brain network organization and its alterations in SZ. An overview of our approach is presented in Figure 1.

**Figure 1:**
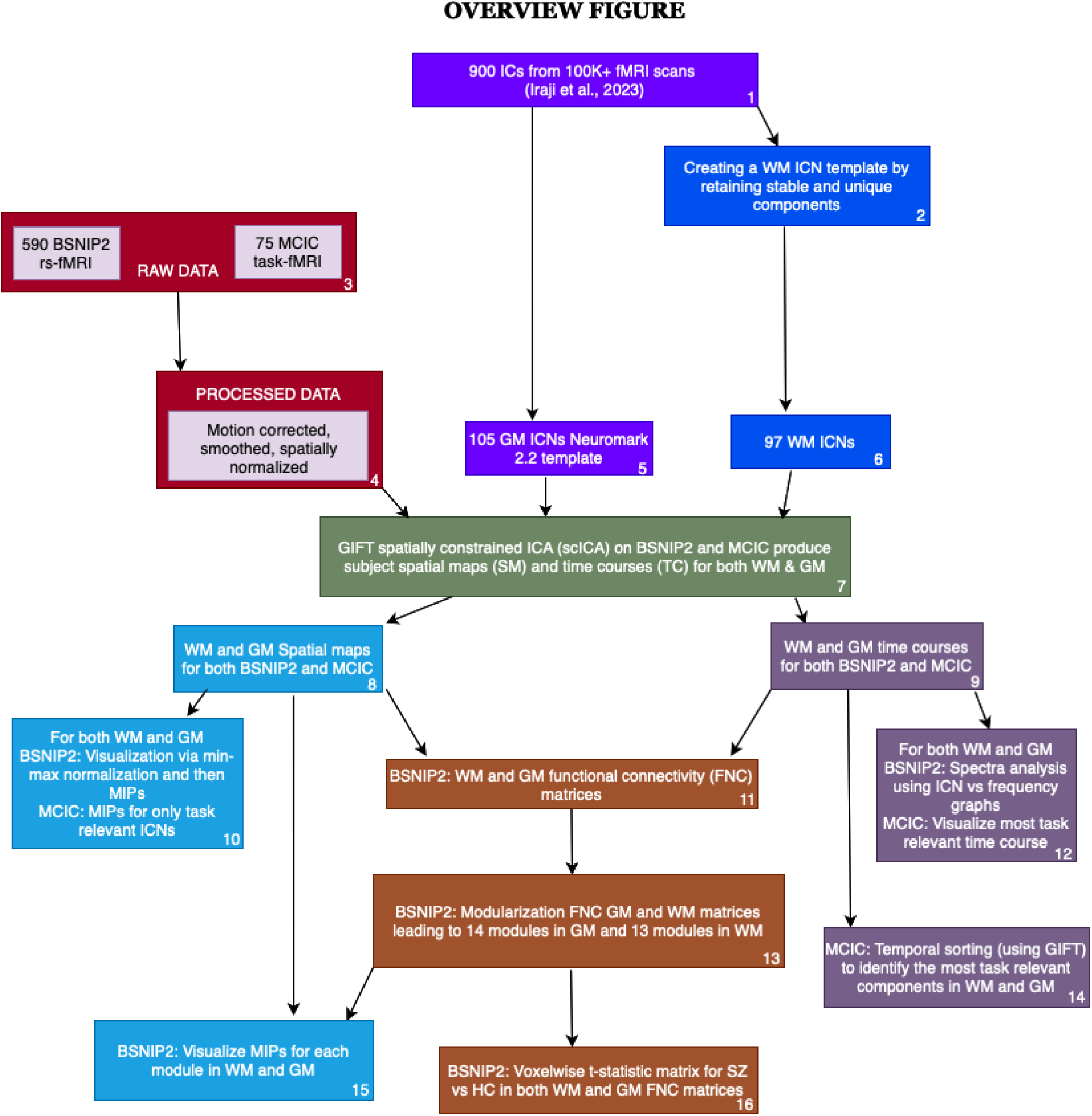
Overview of our approach. The boxes numbered 1, 2, 5, and 6 indicate an open-source dataset (i.e. 900 ICs as well as 105 GM ICNs) and they helped us create a new robust and stable WM template with 97 ICNs (i.e., boxes 2, 6). The boxes 3 and 4 indicate the resting and task-based fMRI dataset we used to apply our template on using NeuroMark and GIFT scICA (indicated by box 7). Light blue boxes (8, 10, 15) represent the visualization of spatial maps to understand the summary of ICNs in WM and GM templates as well as after applying ICA to the BSNIP2 and MCIC datasets. We summarized the spatial maps using maximum intensity projections (MIPs) where we first normalized the dataset for each ICN and then computed the maximum along each voxel for the different ICNs, providing a single image which summarizes all the components. Note this was done separately for GM and WM ICNs. Purple boxes (9, 12, 14) indicate the processing that was done with the time courses for both BSNIP2 and MCIC datasets.

We did temporal sorting for the MCIC dataset to identify the most task-relevant ICNs whereas for BSNIP2 we did spectral analysis. Brown boxes (11, 13, 16) summarize our analysis with the sFNC matrices for the BSNIP2 dataset which helped us identify different modules based on connectivity in both WM and GM.

#### 1. Data Summary

##### Template generation

For our study, we used a comprehensive template for ICNs, drawing from an extensive dataset of 100,517 rs-fMRI scans (Iraji et al., 2023). This dataset included more than 20 private and public sources (refer to supplementary information for more details (Iraji et al., 2023)), representing a wide range of demographic characteristics and imaging protocols. Preprocessing steps included rigid body motion correction, slice timing correction, distortion correction, and spatial smoothing (Iraji et al., 2023). We used available preprocessed data when possible and performed additional preprocessing as needed using FSL and SPM12 toolboxes. Data were then warped into Montreal Neurological Institute (MNI) space and resampled to 3 x 3 x 3 mm³ isotropic voxels. After applying stringent quality control (QC) criteria, which required a minimum of 120 time points in the rs-fMRI time series, mean framewise displacement less than 0.25, head motion transition less than 3° rotation and 3 mm translation in any direction, high-quality registration to an echo-planar imaging template (V. D. Calhoun et al., 2017), we selected 57,709 scans for analysis. Using this rigorously curated dataset, we applied group-level multi-model-order spatial ICA to generate a robust and replicable ICN template across multiple spatial scales (Iraji et al., 2022, 2023). This process involved splitting the QC-passed data, running ICA with various model orders, and selecting the most stable ICNs in white matter based on spatial similarity. The final WM ICN template represents a diverse array of brain patterns, validated through comparison with a separate QC-failed dataset to ensure generalizability and robustness. This high-quality ICN template, derived from a large and diverse dataset, provides a reliable framework for investigating brain connectivity patterns, supporting the functional relevance of white matter in cognitive processes, and paving the way for further research into neural dynamics and disease models.

##### Preprocessing of the fMRI data

Prior to analysis, we preprocessed the BSNIP2 and the MCIC dataset using a robust pipeline. Preprocessing steps included rigid body motion correction, slice timing correction, distortion correction, and spatial smoothing (Iraji et al., 2023). Both datasets were preprocessed using Statistical Parametric Mapping 12 (https://www.fil.ion.ucl.ac.uk/spm/software/spm12/) in Matlab. The first ten scans were discarded to ensure the retention of scans reflecting a stable scanner state. Images were realigned using INRIalign—a motion correction algorithm unbiased by local signal changes (Freire et al., 2002; Freire & Mangin, 2001). A slice-timing correction was performed on the fMRI data after realignment to account for possible errors related to the temporal variability in the acquisition of the fMRI datasets. Data were spatially normalized (Ashburner & Friston, 1999, 2005) into the standard Montreal Neurological Institute (http://www.mni.mcgill.ca/) space using an echo-planar imaging (EPI) template (V. D. Calhoun et al., 2017). The resting-state fMRI data was smoothed with a 6mm full width at half maximum (FWHM) Gaussian kernel and the task data was smoothed slightly more at 9mm due to the smaller number of subjects (Mikl et al., 2008).

##### Task data (MCIC)

We used 75 scans from the sensorimotor task dataset from MCIC’s (Gollub et al., 2013) University of New Mexico (UNM) site. The MCIC schizophrenic dataset used is a publicly accessible, online repository containing curated anatomical and functional MRI. Individuals with SZ and controls were demographically matched, by sex and age. The data can be obtained through the COllaborative Informatics Neuroimaging Suite (COINS; http://coins.trendscenter.org). We used the sensorimotor task fMRI data from this dataset for our analysis. Participants were aged between 18 and 60 and were native English speakers. Inclusion criteria for the SZ cohort required patients to meet DSM-IV diagnostic criteria (Mittal & Walker, 2011) for schizophrenia. fMRI scans involved prospective acquisition correction (PACE) (van der Kouwe et al., 2006). Whole-brain, single-shot EPI data were acquired parallel to the AC-PC line with TR of 2 seconds, TE of 30 milliseconds, flip angle of 90°, in plane resolution of 3.4 mm and 27 slices with a thickness of 4 mm and a 1 mm skip, interleaved slice order. For the sensorimotor task, the MCIC investigators (Gollub et al., 2013) designed a paradigm to robustly activate the auditory and somatosensory cortices by having subjects, with eyes closed, respond to a series of binaural audio tones of varying frequencies using unilateral button presses. Auditory stimuli, delivered through sound-insulated earphones, were calibrated for each subject, and presented in ascending and descending sequences during 16-second auditory blocks alternated with fixation blocks. QC criteria were the same as mentioned above.

##### Rest-state fMRI data (BSNIP2)

For our analysis, we used the BSNIP2 (Tamminga et al., 2014) data which included 590 resting-state functional magnetic resonance imaging (rs-fMRI) scans, with 252 SZ and 338 HC. The BSNIP population consists of research participants gathered from six sites across the United States of America. This sample includes individuals diagnosed with SZ and HC with no immediate family history of psychotic disorders. All testing materials and procedures were standardized and uniformly implemented across all sites. Consensus diagnoses according to DSM-IV criteria (Mittal & Walker, 2011) were determined by trained clinical raters and senior diagnosticians using comprehensive clinical data and structured clinical interviews (SCID) (Spitzer et al., 1992). The inter-rater reliability among the raters was greater than 0.90. Functional scans were obtained with gradient-echo echo planar imaging, featuring a repetition time (TR) of 1.5 seconds, an echo time (TE) of 28 milliseconds, a flip angle of 65°, a voxel size of 3.4 mm × 3.4 mm × 5 mm, a slice thickness of 5 mm, and 30 slices per scan. To reduce head movement, a custom-built head coil cushion was employed. During the scan, subjects were instructed to fixate on a cross on the monitor, stay alert with their eyes open, and keep their head still. QC criteria were the same as mentioned above.

#### 2. Creating the WM Template

We followed an approach similar to (Iraji et al., 2023) which introduced a multi-scale gray matter template. In our case, to generate the WM template, we use the set of 900 independent components generated from the multi-scale ICA (Iraji et al., 2023) to identify the subset of replicable WM ICNs. Our goal was to extract a WM ICN template with robust stability, similar to the process for deriving the existing GM template comprising 105 ICNs (Iraji et al., 2023). Our process first involved curation, starting with the elimination of ICNs lacking maximum presence in WM regions and those exhibiting instability (stability < 0.8). Subsequently, we pruned duplicate ICNs by identifying them through high similarity scores (*r* > 0.8). This iterative pruning process left us with approximately 110 putative WM ICNs. To ascertain the predominant presence of the WM ICN in the WM tract, we performed additional manual scrutiny by experts who are familiar with WM structures, cross-referencing them with established GM and WM templates as well as the Automated Anatomical Atlas (AAL, (Tzourio-Mazoyer et al., 2002)) GM atlas and JHU WM atlas (Mori et al., 2005). This meticulous validation process yielded a final selection of 97 WM ICNs that demonstrated stability and uniqueness, named NeuroMark WM template, thereby forming the foundation of our WM template.

To visualize the spatial distribution of ICNs in WM and GM, we employed MIPs using WM and GM ICN template data. MIPs method involved normalizing each ICN using minimum-maximum normalization and then computing MIPs i.e., by computing the maximum value along each voxel axis for spatial maps derived from independent ICNs, facilitating the visualization and comparison of functional brain networks. The resulting MIPs for the 105 GM and 97 WM ICNs were thresholded at 0.65 to highlight the most significant regions. To further examine the relationship between WM ICNs and specific tracts, we determined the maximum voxel intensity and matched each WM ICN with the resampled JHU white matter atlas. MIPs were then generated for each corresponding WM tract, providing a clearer visualization of their anatomical relevance.

#### 3. GIFT scICA and sFNC on templates

Our study employed scICA from the GIFT toolbox (https://trendscenter.org/software/gift/) (V. d. Calhoun et al., 2001; Iraji et al., 2021) to analyze the BSNIP2 and MCIC datasets, focusing on both WM and GM functional connectivity. Using our hybrid NeuroMark pipeline which combined scICA with replicable spatial priors derived from large independent datasets from more than 100,000 scans (Iraji et al., 2023). These spatial priors include 105 GM ICNs from multiple spatial scales which have been previously organized and described (Jensen et al., 2024) and are freely available in the Neuromark_fMRI_2.2_modelorder-multi in GIFT (also available separately at https://trendscenter.org/data/). The current study extends this GM template by introducing an additional 97 WM ICNs. The WM ICNs were combined with the GM template (105 GM ICNs; Iraji et al., 2023; Jensen et al., 2024) for our current investigation. These two templates provided foundational ICNs for subsequent analyses, facilitating the characterization of functional connectivity patterns within both GM and WM regions.

#### 4. Analysis and comparison of GM and WM spatial maps, time courses, and sFNC

To improve the accessibility of the NeuroMark WM template, we implemented several key modifications: First, we visually inspected each of the 97 WM ICNs and made notes on their spatial overlap with white matter in the JHU WM atlas (Mori et al., 2005). Next, we classified these WM ICNs into domains and subdomains, using standard neuroscience terminology typically used to describe GM regions. Finally, we reorganized the ICNs based on the sFNC using both GM and WM matrix combined, grouping WM ICNs in a domain comparable to the GM domains based on the GM ICNs with the highest correlation, which improved the functional interpretability of the ICNs. These enhancements aim to facilitate the usability of the WM ICN NeuroMark template within the neuroscience community. Each of the 97 WM ICNs was thoroughly examined using MRIcroGL (Rorden & Brett, 2000) available at https://www.nitrc.org/projects/mricrogl, where they were overlaid on the JHU white matter atlas and cross-referenced with the AAL atlas. To provide additional anatomical context, we identified the nearest gray matter (GM) regions using the AAL atlas. We calculated the spatial similarity between the WM ICNs and their corresponding GM regions, documenting details such as peak voxel locations, size, and shape for each ICN. Furthermore, we modified existing subdomains to capture the distinctive features of the WM ICNs. For instance, WM ICNs surrounding the GM sensorimotor domain were categorized as posterior, middle, or anterior sensorimotor based on their location relative to the GM sensorimotor domain. This division of the GM sensorimotor domain was done to help distinguish between the large number of WM ICNs in these regions of the brain, which are likely a result of the large number of WM tracts in these regions. This structured approach provides a clearer understanding of the spatial arrangement of WM ICNs within white matter pathways, thereby enhancing the interpretability of the WM template. The resulting domain, subdomain, and individual labels have been released with WM ICN NeuroMark template.

After extracting time courses and spatial components for each subject, we conducted temporal filtering and removed participants with high motion artifacts. The sFNC was calculated following standard preprocessing steps, Pearson correlations were computed between the time courses of all 202 subject-specific ICNs, comprising 97 WM ICNs and 105 GM ICNs, for each individual (Du et al., 2020). These correlation values were transformed into z-scores and then averaged across subjects to obtain a group-level sFNC matrix as shown in Figure 6. The resulting sFNC matrix reflects inter-network connectivity by representing the strength of interaction between different ICNs. While the spatial map of each ICN illustrates intra-network connectivity, the sFNC matrix highlights the connectivity strength between distinct networks.

After having the spatial maps of ICNs, their corresponding TCs, and the sFNC between TC, we conducted a multifaceted analysis encompassing several key steps. Primarily, we computed the MIPs of the spatial maps of the ICNs using both WM and GM ICN templates, seperately. To understand more about the frequency profile of the WM and GM ICNs, the filtered time series underwent transformation into the frequency domain using a fast Fourier transform to obtain the power spectrum. Subsequently, we analyzed the temporal characteristics of these networks, generating frequency graphs and summarizing power spectra to elucidate intrinsic oscillatory patterns. sFNC analysis was performed on 590 subjects from the BSNIP2 dataset, providing insights into brain network organization and its modulation.

#### 5. Task-Based fMRI Analysis

In our task-based fMRI analysis, we adopted a comprehensive approach to examine the neural correlates of cognitive tasks. We began by performing scICA on both smoothed and unsmoothed MCIC data to ensure the independence of WM ICNs from GM ICNs. The comparison between smoothed and unsmoothed data was motivated by the need to evaluate whether smoothing, which enhances signal-to-noise ratio but may blur fine-grained boundaries, impacts the independence of WM and GM ICNs, thereby ensuring robust and consistent findings across preprocessing approaches. Following this, we conducted temporal sorting, a process that aligns the time course of ICNs with external task-related timing information, was conducted to assess task-related effects, calculating metrics such as R^2^ and beta values using the GIFT toolbox. From the results of the temporal sorting, we identified the two ICNs in both WM and GM that showed the strongest associations with the task paradigm. Specifically, we selected the most relevant ICNs based on significant p-values (p < 0.001) and their corresponding R^2^ values (see Figure 9A). We also computed the MIPs for all significant ICNs (p < 0.001) in both GM and WM are illustrated in Figure 10.

We established the onsets and durations of the stimulus, which were essential for time-locking our data. Each block comprised a 1 TR off period, followed by an 8 TR active phase, and concluded with a 9 TR off period, resulting in a total block length of 18 TR. To compute time-locked averages for the top two task significant GM and WM ICNs across blocks, we applied a baseline shift method. This involved subtracting the minimum value of each segment from the segment itself, thereby normalizing the data. Additionally, we performed interpolation to enhance the resolution of our time-locked averages, allowing for more precise visualizations. The interpolation was executed with an interpolation factor of 10, effectively increasing the temporal resolution of the signal by interpolating each segment to obtain a finer time course. For the GM and WM time components, we computed the time-locked averages after interpolation. We subsequently plotted the mean signals of the GM and WM ICNs against time in milliseconds. This allowed us to visualize the relationship between WM and GM activations over time.

#### 6. Group Difference Analysis

We analyzed functional connectivity patterns using group-specific sFNC matrices for subjects with SZ (n = 252) and HC (n = 338). To identify aberrant connectivity patterns associated with SZ, we performed a Generalized Linear Model (GLM) analysis. The model accounted for confounding variables, including age, sex, race, site, and head motion. This approach allowed us to calculate voxel-wise t-statistics and p-values for each FNC pair, with statistical significance determined using false discovery rate (FDR) correction (q < 0.05). The resulting t-statistic values were visualized in a connectivity map, highlighting patterns of altered functional connectivity in SZ (see Figure 11).

## Results

### 1. Visualization of the GM and WM ICN Templates

The resulting MIP images of the 105 GM ICN and 97 WM ICN templates are depicted in Figure 2, thresholded at 0.65, revealing distinct regions anatomically aligned with the GM and WM.

**Figure 2:**
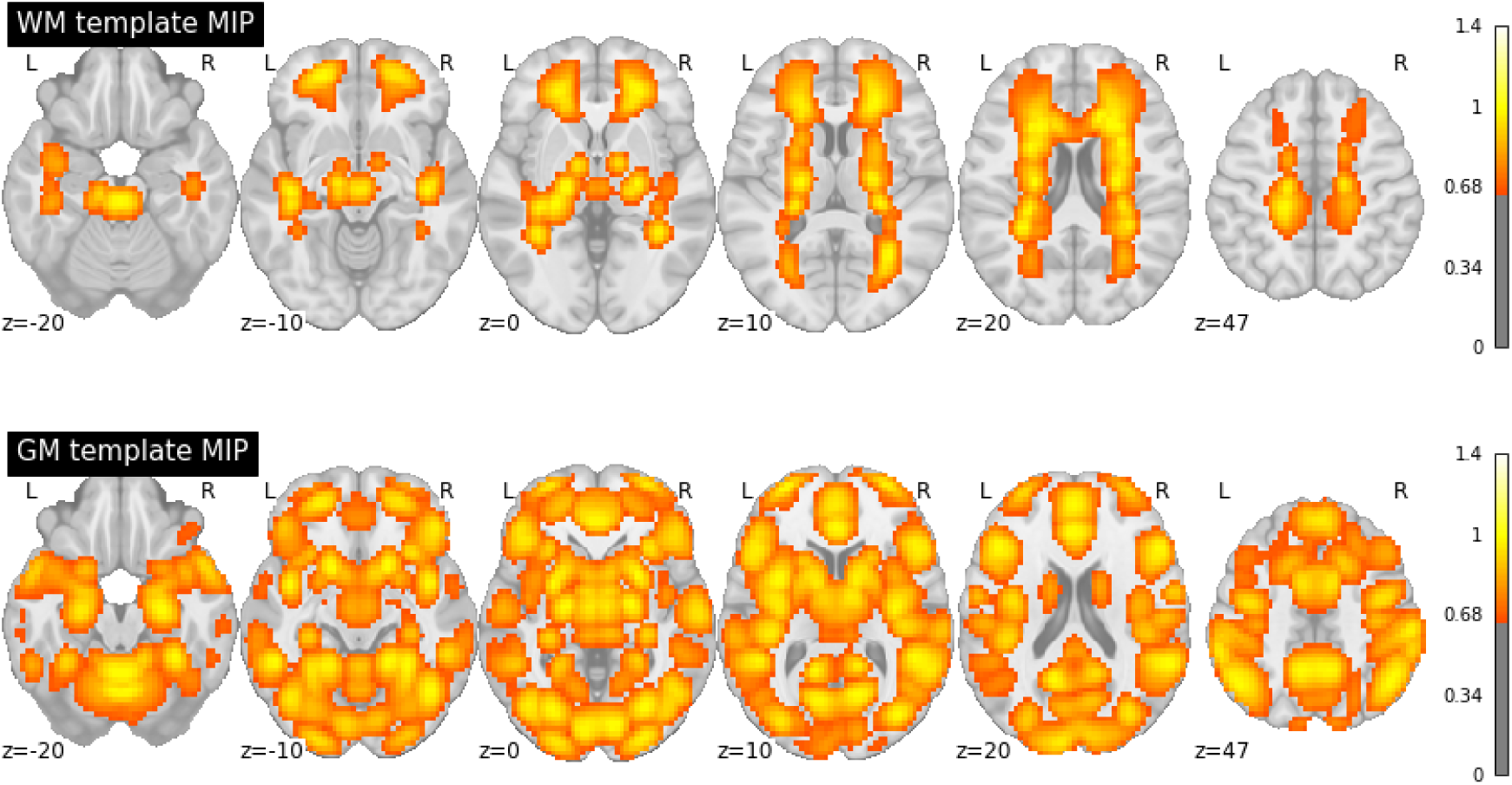
MIPs of GM and WM templates (smoothed) derived from the large sample dataset. We could distinctly differentiate anatomically aligned regions of GM and WM, highlighting the clear separation between the 105 GM ICN and 97 WM ICN templates.

The resulting MIPs (Figure 3) from the tract-based allocation of each WM ICN demonstrated that our WM ICN template effectively extracted both unilateral and bilateral WM components. We successfully classified the 97 WM ICNs into 24 distinct WM tracts using the JHU atlas. The tracts included in our template are the anterior corona radiata (r,l), the retrolenticular part of internal capsule (r,l), the superior corona radiata (r,l), the middle cerebellar peduncle, the posterior corona radiata (r,l), the cingulum, the posterior limb of internal capsule (r,l), the pontine crossing tract, the cerebral peduncle (l), the posterior thalamic radiation, the urcinate fasciculus (l), the body of corpus callosum, the sagittal stratum, the anterior limb of internal capsule (r,l), the superior longitudinal fasciculus (r,l), the tapetum (r) and some unclassified tracts. This classification indicates the template’s capability to accurately map WM pathways and provides a clear representation of the spatial organization of WM ICNs across the brain.

**Figure 3:**
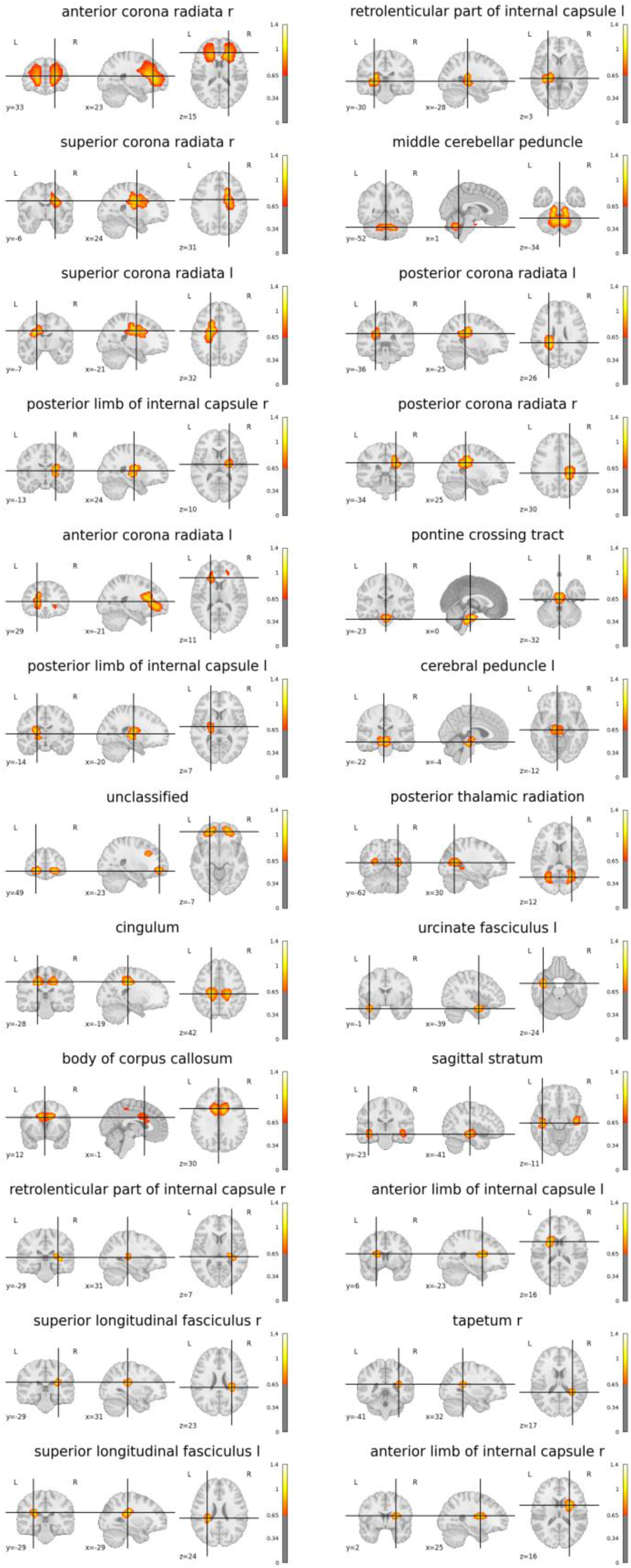
MIP maps (smoothed) summarizing the components falling within each of the WM tracts in the 97 WM ICN template.

**Figure 4:**
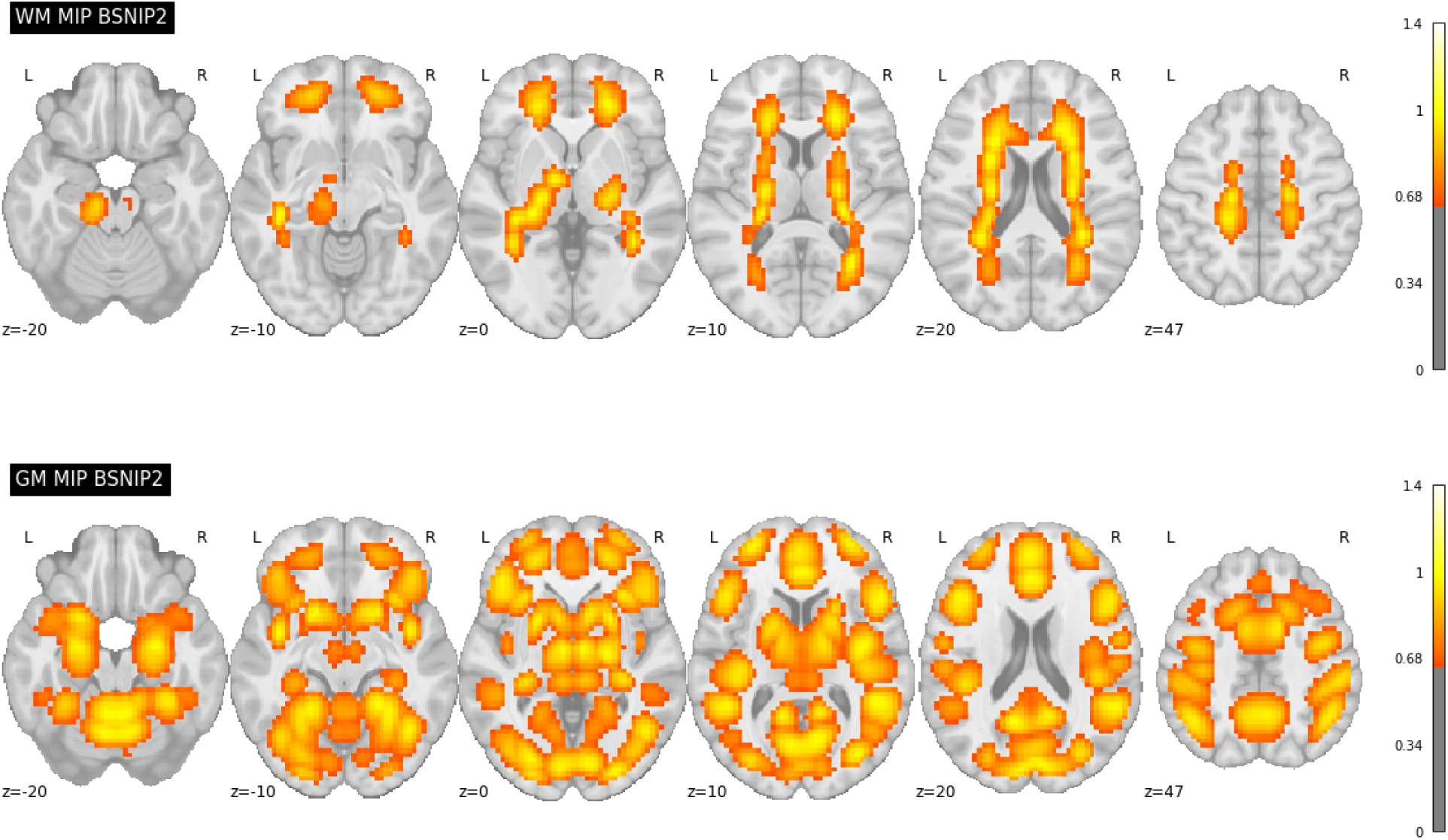
MIPs of smoothed GM and WM ICNs derived from the BSNIP2 scICA output. The distinct functional connectivity patterns between GM and WM are evident, further reinforcing the inherent differences between these tissue types.

### 2. Resting-State fMRI Result

We further examined the BSNIP2 rs-fMRI data by generating MIP images from the BSNIP2 scICA output for GM and WM ICNs as shown in 4. As expected, these maps also show distinct WM/GM peak locations but there are also clear differences from Figure 2. This reflects the individual subject and dataset variability which is captured by the scICA approach.

#### 2.1 Frequency Analysis of ICNs of BSNIP2

An additional higher frequency band in WM ICNs was observed as shown in Figure 5A by comparing averaged power spectra for both GM and WM ICNs. This discovery prompted a deeper investigation into the spectral properties of these networks. By summarizing the frequency profiles using a TC spectrum and averaging across components, we plotted a line graph as shown in Figure 5B that confirmed the presence of this additional spectral peak around 0.06 Hz in WM ICNs, as well as slightly more power at the higher frequencies, distinguishing it further from the GM ICNs.

**Figure 5:**
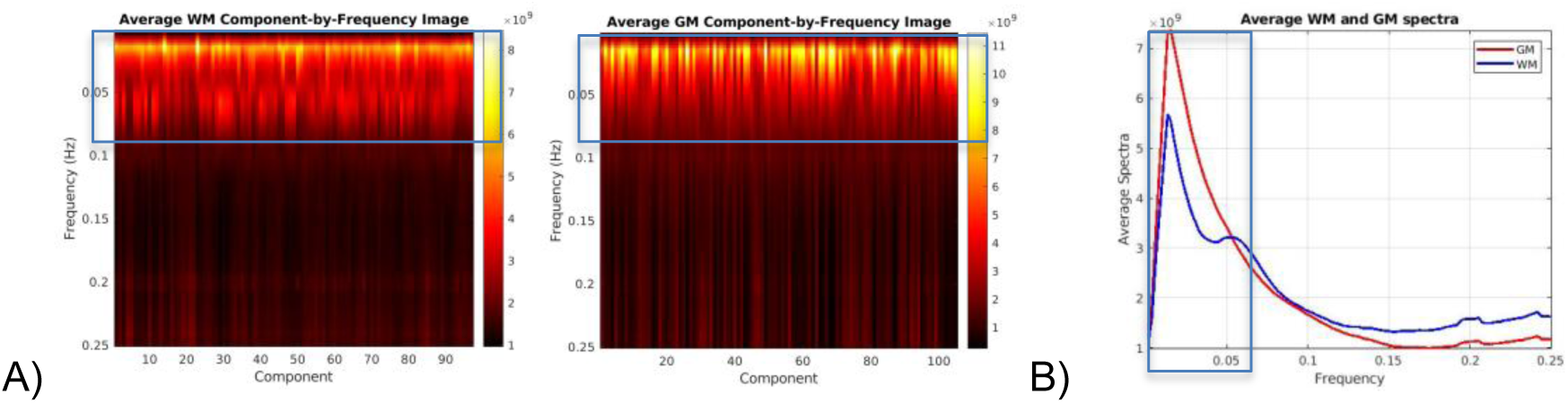
Frequency spectrum analysis of WM and GM ICNs. A higher frequency peak around 0.06 Hz is observed in WM ICNs, distinguishing their spectral properties from GM ICNs. A) Comparing average ICNs vs. frequency graphs for both GM and WM ICNs. B) To inspect the high-frequency band in WM (indicated in the box), this panel shows power spectrum after averaging across ICNs.

#### 2.2 GM and WM sFNC modularization using BSNIP2

The collective and ordered representation of the entire WM and GM sFNC matrix (Figure 6) allows us to discern overarching patterns of functional connectivity within the studied population, aiding our understanding of the brain’s intrinsic network dynamics. The GM sFNC matrix was modularized based on the NeuroMark 2.2 template (Jensen et al., 2024), resulting in the following 14 distinct modules: cerebellar (CB), visual-occipitotemporal (VI-OT), visual-occipital (VI-OC), paralimbic (PL), subcortical-extended hippocampal (SC-EH), subcortical-extended thalamic (SC-ET), subcortical-basal ganglia (SC-BG), sensorimotor (SM), higher cognition-insular temporal (HC-IT), higher cognition-temporoparietal (HC-TP), higher cognition-frontal (HC-FR), triple network-central executive (TN-CE), triple network-default mode (TN-DM), and triple network-salience (TN-SA). The WM sFNC matrix was modularized, resulting in the following 13 modules: paralimbic (PL), subcortical-posterior hippocampal (SC-PH), subcortical-thalamic hippocampal (SC-TH), subcortical-extended thalamic (SC-ET), subcortical-basal ganglia (SC-BG), frontal (FR), sensorimotor-middle sensorimotor (SM-MS), sensorimotor-anterior sensorimotor (SM-AS), sensorimotor-posterior sensorimotor (SM-PS), insular (IN), temporoparietal (TP), occipitotemporal (OT), cerebellar-brainstem (CB; see Figure 7). High modularity can be observed within subdomains (the modules represent subdomains) as well as between spatially proximal WM and GM subdomains (e.g., between the subcortical WM and subcortical GM subdomains). Our analyses revealed that high functional connectivity (FC) between WM ICNs is not simply based on spatial proximity. Instead, we observed high FC between ICNs along the same WM tract with little to no spatial overlap. To investigate whether these observed patterns of modularity were due to functional similarity or merely an effect of spatial smoothing, we identified two WM ICNs from different subdomains which were located along the same WM tract (i.e., the corona radiata) and observed that they elicited a relatively high correlation (0.4954) despite having little or no spatial overlap (see Figure 8). This observation provides evidence of a unique functional signature in WM,, reinforcing the idea that WM networks actively contribute to brain function rather than merely providing structural support.

**Figure 6:**
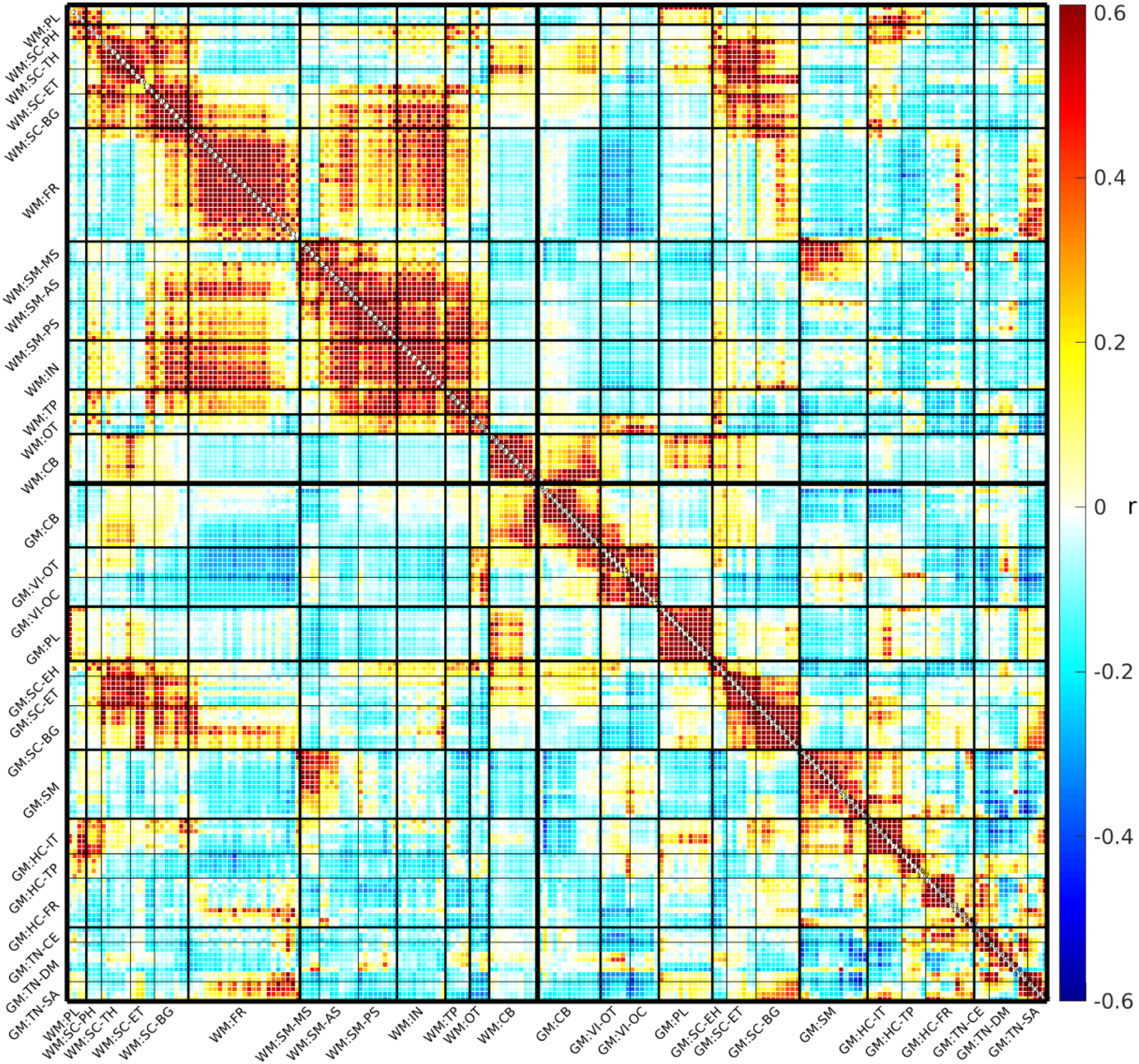
Group-level averaged sFNC matrix. The sFNC matrix is modularized into 13 white matter (WM) subdomains and 14 gray matter (GM) subdomains, indicated with a ‘WM:’ or ‘GM:’ prefix respectively. The 13 WM modules are: paralimbic (PL), subcortical-posterior hippocampal (SC-PH), subcortical-thalamic hippocampal (SC-TH), subcortical-extended thalamic (SC-ET), subcortical-basal ganglia (SC-BG), frontal (FR), sensorimotor-middle sensorimotor (SM-MS), sensorimotor-anterior sensorimotor (SM-AS), sensorimotor-posterior sensorimotor (SM-PS), insular (IN), temporoparietal (TP), occipitotemporal (OT), cerebellar-brainstem (CB). The following are the 14 GM modules: cerebellar (CB), visual-occipitotemporal (VI-OT), visual-occipital (VI-OC), paralimbic (PL), subcortical-extended hippocampal (SC-EH), subcortical-extended thalamic (SC-ET), subcortical-basal ganglia (SC-BG), sensorimotor (SM), higher cognition-insular temporal (HC-IT), higher cognition-temporoparietal (HC-TP), higher cognition-frontal (HC-FR), triple network-central executive (TN-CE), triple network-default mode (TN-DM), and triple network-salience (TN-SA). For spatial maps and additional information, see Figure 7 and supplementary materials for the WM modules, and see Jensen et al. (2024) for the GM modules.

**Figure 7:**
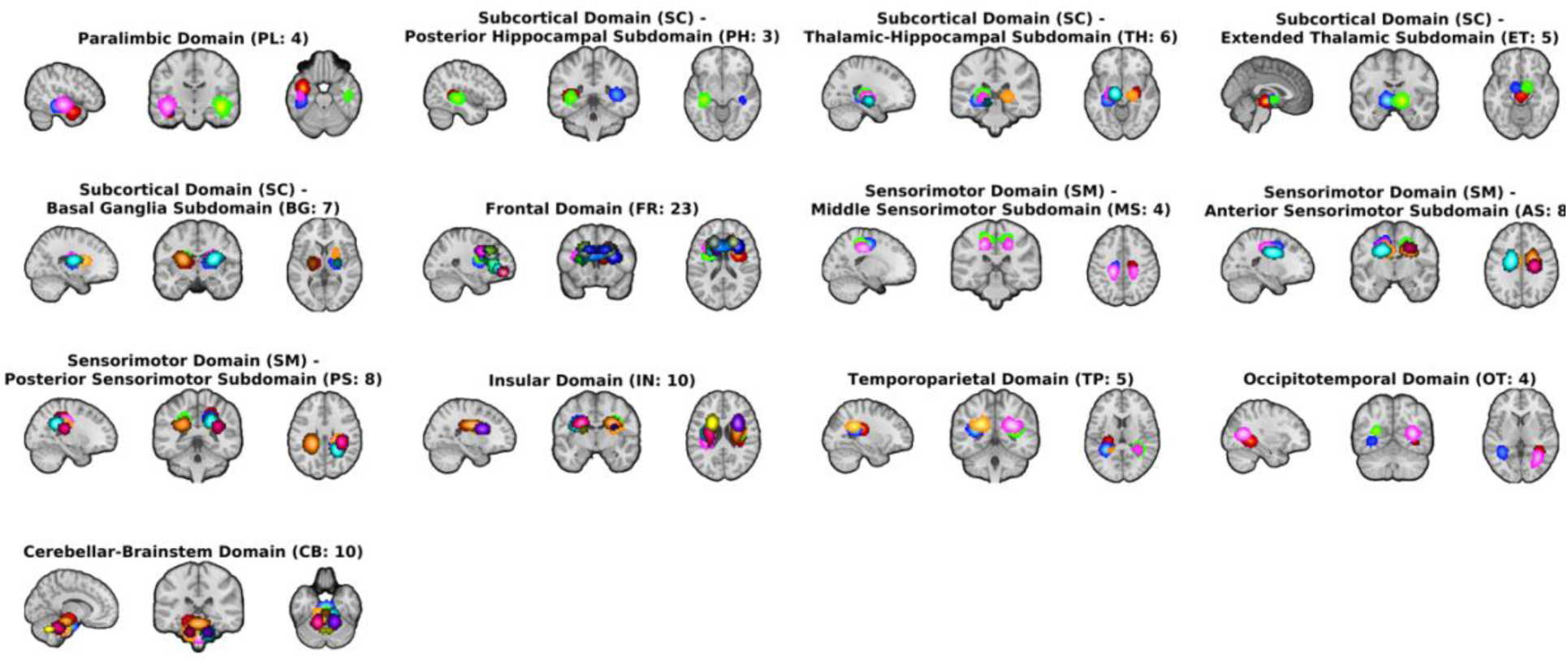
Visualization of the domains and subdomains (modules) for the WM ICNs using MIPs.

**Figure 8:**
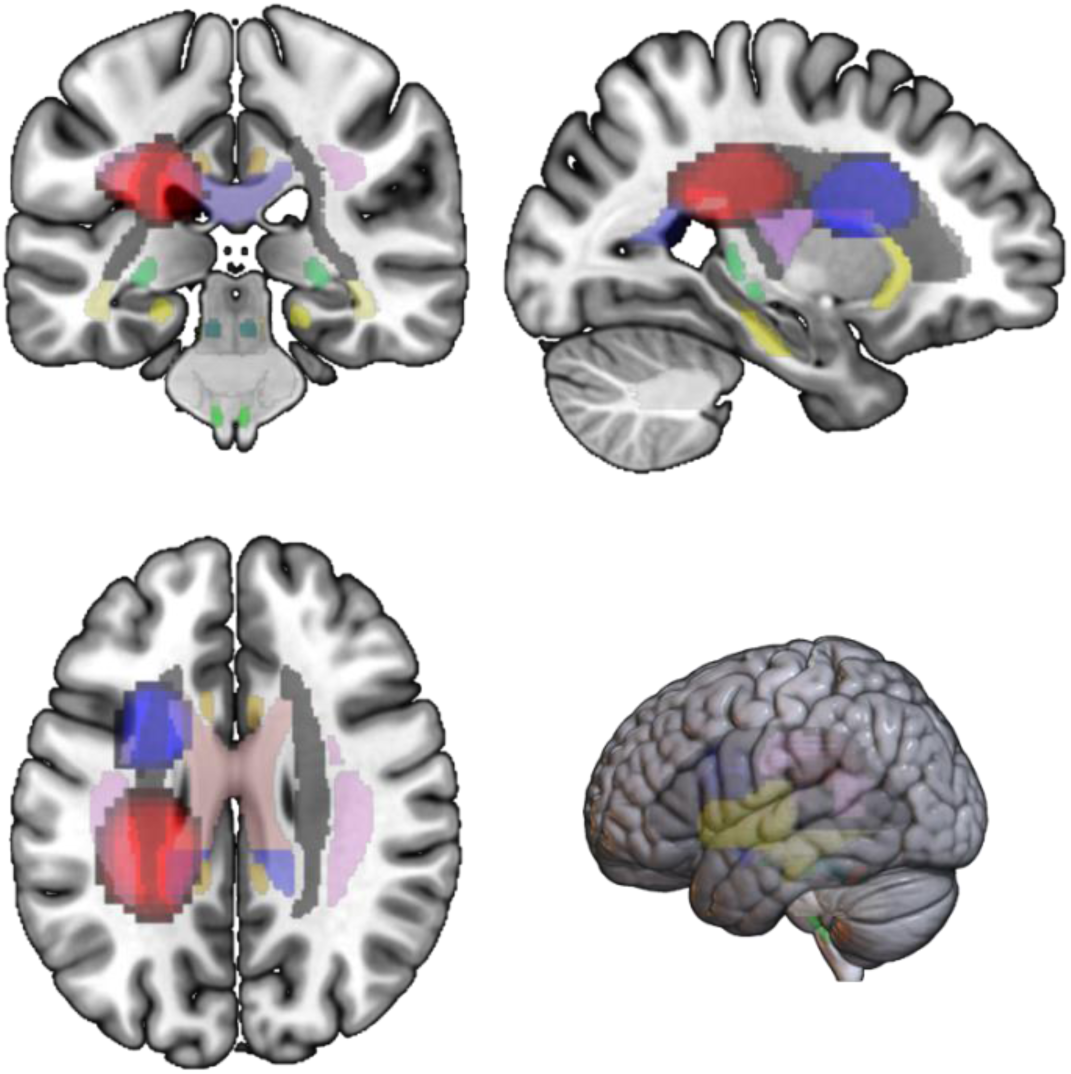
High correlation between two WM ICNs overlapping with the corona radiata. Depicted above are WM ICN 67 in red and WM ICN 28 in blue. WM ICN 67 spatially overlaps with the left posterior corona radiata and is grouped into the posterior sensorimotor domain, whereas WM ICN 28 spatially overlaps with the left superior and anterior corona radiata and is grouped into the frontal domain. Both of these WM ICNs have been overlayed on the MNI152 and JHU atlas with a threshold set between a z-score of 5 and 9. The correlation value between these two ICNs (shown in Figure 6) is 0.50 and the coordinates for the slice displayed above are as follows: - 26.35 mm (x), -31.17 mm (y), 28.53 mm (z).

### 3. Task-Based fMRI Results using MCIC

The time-locked averages in Figure 9B, reveal that WM TC has an earlier rise after stimulus compared to GM TC. From Figure 9A, we observe that GM ICN 75 is associated with the bilateral auditory region, while GM ICN 62 likely corresponds to the left hemisphere sensorimotor area, specifically linked to finger tapping. This aligns with the task-based fMRI data, which involved bilateral auditory stimulation followed by a sensorimotor task (tapping with the dominant hand’s finger). The TC in Figure 9B and spatial maps in Figure 9A further show that the top two WM ICNs i.e., WM ICN 7 and WM ICN 5 align with the superior longitudinal fasciculus, a white matter tract crucial for integrating auditory and sensorimotor functions. Interestingly, we observed distinct motor and auditory ICNs, as expected for this task. The corticospinal tract also displayed lateralization consistent with motor movements on the same side as the motor ICNs, reinforcing the role of WM in motor activity. Notably, WM TC signals decayed before GM indicating that WM TC attains a post-stimulus minimum before the GM TC, suggesting to further explore this phenomenon during task-related processing.

**Figure 9:**
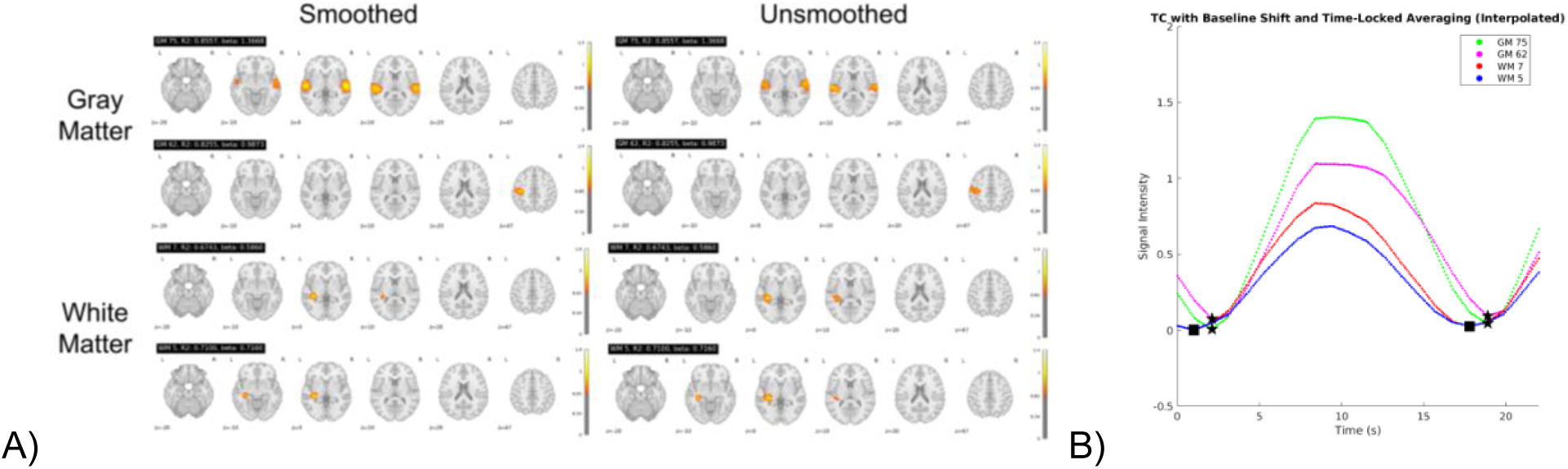
Temporal Sorting and Activation Patterns in Task-Based fMRI. A) Visualization shows distinct motor and auditory components. B) Time courses with baseline shift and time-locked average showing GM and WM signals rising after stimulus as well as a post stimulus decay. Here, star represents the GM TC minimum whereas square represents the WM TC minimum for two situations i.e., rise after stimulus (before 200 ms) and post stimulus minimum (after 800 ms).

To validate that these WM activations were not merely extensions or blurs of GM changes, we compared smoothed and unsmoothed spatial ICN maps (Figure 9A and Figure 10). The results confirmed that WM activations were distinct and did not appear to be artifacts of GM signal spillover. Figure 10 summarizes the MIPs of all the task-significant (p<0.001) and task insignificant (p>0.001) WM and GM ICNs.

**Figure 10:**
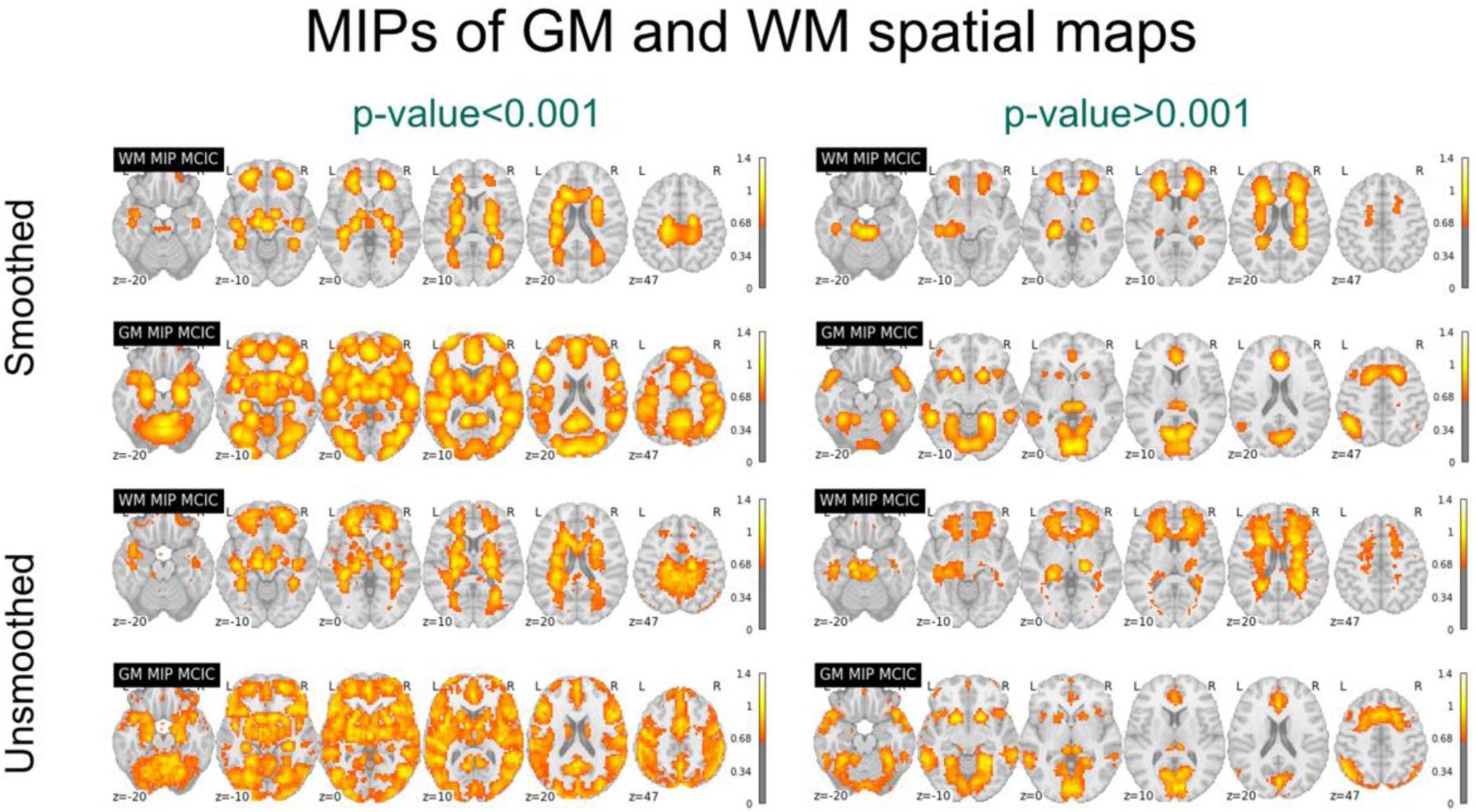
Comparing significant (p<0.001) and insignificant task-relevant MIPs in the MCIC dataset. This proves that task-relevant activation in WM is different from GM and not affected due to the smoothing effects. Task-relevant ICNs were identified in both GM and WM using significant p-values, indicating the engagement of both tissue types in cognitive tasks.

### 4. Group differences (SZ vs HC) in BSNIP2

The t-statistic matrix (Figure 11) illustrates significant group-level differences in functional connectivity patterns between SZ patients and HC individuals, revealing distinct alterations across both GM and WM domains. The sFNC t-value matrix was derived using a GLM model, shows striking results: blue regions indicate negative t-values (SZ<HC), while red regions represent positive t-values (SZ>HC). In the WM sFNC matrix, a substantial amount of blue suggests a marked reduction in functional connectivity in SZ patients compared to HC, particularly in the second module. This reduced connectivity, observed in regions such as the insular, temporoparietal, and subcortical extended thalamus, may reflect disruptions in white matter pathways crucial for efficient communication between brain regions. Additionally, we notice significant inter-WM domain differences, with decreased connectivity between the frontal and subcortical basal ganglia, frontal and temporoparietal, frontal and insular, and frontal and sensorimotor ICNs, suggesting a widespread decline in cross-domain WM connectivity in SZ. Conversely, the GM matrix presents a pattern inclusive of both red and blue regions, indicating areas of potential hyperconnectivity (red) in SZ, such as in the sensory-motor and SC-ET (subcortical-extended thalamus) regions, where SZ patients exhibit greater connectivity than HC. This heightened activity in certain GM regions might reflect compensatory mechanisms or altered functional processing. The combined GM and WM sFNC matrix, particularly in the first and third quadrants, reveals further intriguing differences, with several red regions indicating that SZ patients exhibit increased connectivity between WM frontal and GM regions related to higher cognition, insular, and temporal areas compared to HC. This complex interaction between GM and WM networks underscores the broader dysfunction in brain connectivity in SZ, where WM disruptions and GM hyperactivity may jointly contribute to the neurobiological basis of the disorder.

**Figure 11:**
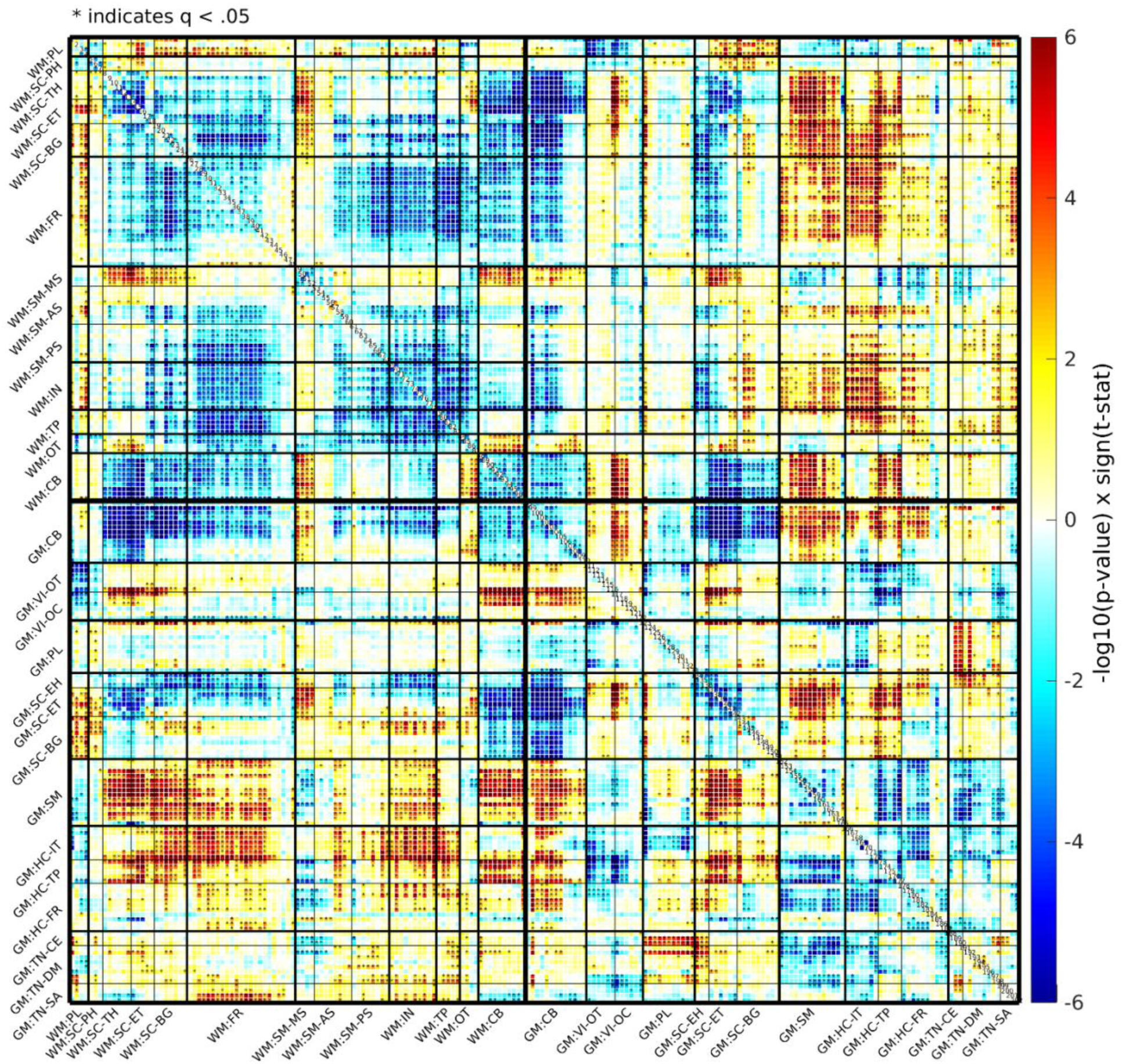
White matter (WM) and gray matter (GM) Connectivity Differences Between SZ and HC Groups in BSNIP2 (SZ < HC in blue, SZ > HC in red). T-statistic matrix revealing differences in functional connectivity patterns between SZ patients and HC, particularly in the second module. The labels with a prefix ‘WM:’ indicate the WM modules whereas the remaining ones with a prefix ‘GM:’ indicate the GM modules from the Neuromark 2.2 template. The 13 WM modules are: paralimbic (PL), subcortical-posterior hippocampal (SC-PH), subcortical-thalamic hippocampal (SC-TH), subcortical-extended thalamic (SC-ET), subcortical-basal ganglia (SC-BG), frontal (FR), sensorimotor-middle sensorimotor (SM-MS), sensorimotor-anterior sensorimotor (SM-AS), sensorimotor-posterior sensorimotor (SM-PS), insular (IN), temporoparietal (TP), occipitotemporal (OT), cerebellar-brainstem (CB). The following are the 14 GM modules: cerebellar (CB), visual-occipitotemporal (VI-OT), visual-occipital (VI-OC), paralimbic (PL), subcortical-extended hippocampal (SC-EH), subcortical-extended thalamic (SC-ET), subcortical-basal ganglia (SC-BG), sensorimotor (SM), higher cognition-insular temporal (HC-IT), higher cognition-temporoparietal (HC-TP), higher cognition-frontal (HC-FR), triple network-central executive (TN-CE), triple network-default mode (TN-DM), and triple network-salience (TN-SA).

## Discussion

This study presents an extensive examination of WM ICNs, introducing a novel template derived from a large-scale dataset. Our findings confirm the existence of 97 unique WM ICNs, each exhibiting distinct spatial maps, time spectra and functional connectivity patterns compared to the 105 GM ICNs (Iraji et al., 2023; Jensen et al., 2024). Task-based fMRI analysis demonstrated that WM ICNs are involved in task-related activations, aligning with the behavior of GM ICNs, and underscoring the relevance of WM in cognitive processes. We observed notable differences in sFNC patterns between SZ patients and HC, emphasizing the importance of investigating WM in addition to GM. This discussion delves into the key takeaways, explores the potential applications of the WM template, and outlines exciting avenues for future research.

### Unveiling the Power of the WM Template: A Call for Open-Source Integration

This study introduces a novel WM ICN template derived from a large-scale dataset, serving as a valuable resource for the scientific community. The availability of this template and its potential applications in future research are significant, as it can be integrated into widely used neuroimaging software, facilitating broader studies on WM connectivity (Forkel et al., 2022; Iraji et al., 2023). For instance, our template could be used with the auto-labeler software (Salman et al., 2022) to label new networks in future studies using blind ICA. Alternatively, our template could be used in combination with the network correspondance toolbox for comparison with widely used whole brain parcellations that are not derived using ICA (Kong et al., 2024). Currently, most neuroimaging software packages focus primarily on GM analysis, and integrating the WM template would enhance the adoption of WM ICN analysis. This would enable researchers to replicate and extend upon the current findings, promoting a deeper understanding of WM connectivity. Furthermore, the open-source nature of this template encourages widespread adoption and collaboration, ultimately fostering advancements in our understanding of WM’s role in brain function. The methodological approach utilizing scICA was pivotal for accurately delineating WM and GM networks, allowing for the precise identification of distinct ICNs. Additionally, the NeuroMark framework, which integrates structural and functional data, further strengthens the robustness and applicability of our findings (V. d. Calhoun et al., 2001; Du et al., 2020; Iraji et al., 2023). The reproducibility and generalizability of the ICA-derived ICNs across diverse populations underscore the utility of the NeuroMark framework in both clinical and research settings, promoting its potential for widespread adoption (Iraji et al., 2023). By enhancing the reproducibility and comparability of WM studies, this template also promotes research into personalized diagnostics and therapeutic interventions (Du et al., 2020; Iraji et al., 2023).

### Distinct Characteristics of WM ICNs: A New Dimension in Brain Connectivity

The study provides additional evidence for the presence of WM ICNs as well as their clear spatial distinction from GM ICNs. Spatial maps revealed distinct anatomical distributions, suggesting unique roles for each in brain function (V. d. Calhoun et al., 2001; Du et al., 2020; Iraji et al., 2023). This distinction is further supported by the modularity displayed in the sFNC matrix (see Figure 6). Notably, the domains and subdomains reflect a more uniform and modular pattern with relatively little variation across domains and subdomains. Despite the similar number of unique ICNs (WM = 97, GM = 105), the WM ICNs were less spatially distributed than the GM ICNs resulting in fewer modules or subdomains (WM = 13, GM = 14). The homogenous modules of WM FNC in contrast to the more heterogenous GM appears to be consistent with the notion that WM networks are less specialized than GM networks (Franco et al., 2013; Gawryluk et al., 2014). Additionally, spectral analysis indicated that WM ICNs have unique frequency profiles in these data, including a higher frequency band around 0.06 Hz not observed in GM ICNs, which may be critical for understanding the functional dynamics of WM (Gore et al., 2019). These findings challenge the traditional view of WM as solely providing structural support and emphasize its active participation in brain networks. Diffusion MRI studies have begun to elucidate the microstructure of WM and its potential role in information processing e.g., (Tang et al., 2017; Hu et al., 2023; Wang et al., 2022, 2023)). The current work extends these findings by demonstrating the existence of distinct WM ICNs with unique spatial distributions, modular structures, and spectral properties compared to GM ICNs. This aligns with observations by Iraji et al. (2023) who identified functionally distinct WM networks using resting-state fMRI. These findings suggest a more active role for WM in brain function, warranting further investigation into its specific contributions.

### Dual Approach: Unveiling the Full Spectrum of WM Connectivity

Our dual approach using both resting-state and task-based fMRI provided complementary insights into WM connectivity. This holistic approach is crucial for understanding the full spectrum of brain connectivity (Biswal et al., 1995; M. H. Lee et al., 2013). While resting-state fMRI reveals intrinsic connectivity patterns, task-based fMRI highlighted task-evoked activations within WM ICNs. The current work demonstrates for the first time that WM ICNs exhibit task-related activation, further emphasizing their role in cognitive processes. Additionally, incorporating diffusion MRI, which offers insights into white matter microstructure, could provide valuable information about the structural underpinnings of WM ICN function (Tang et al., 2017). The presence of task-related BOLD signals in WM ICNs, particularly within the corticospinal tract during a sensorimotor task, underscores the functional relevance of WM in cognitive processes and supports its role beyond mere structural support (Gollub et al., 2013; Gore et al., 2019). These findings open potential avenues for further research into the implications of the corticospinal tract in various disease models.

Our study reveals that WM TCs have an earlier rise after stimulus as well as an earlier minimum post stimulus compared to the GM TC, during task-based fMRI. Moreover, we observe our study linked GM ICN 75 to bilateral auditory processing and GM ICN 62 associated with the left hemisphere’s sensorimotor area, particularly during finger tapping. Neural activity is expected to first emerge in the primary auditory cortex, which processes auditory information. The identified WM ICNs, particularly WM ICN 7 and WM ICN 5, are aligned with the superior longitudinal fasciculus, highlighting their role in integrating auditory input and motor output. This underscores the importance of WM pathways in enabling efficient communication between sensory and motor regions during complex tasks. The observed lateralization of the corticospinal tract aligns with motor movements, reinforcing WM’s active participation in task-related activities. The distinct patterns of motor and auditory components reflect the complexity of the neural circuits engaged during task performance, supporting the notion that WM actively contributes to neural computations rather than serving merely as a passive conduit. The clear differentiation between WM and GM activations calls for further investigation into WM’s role in cognitive and sensorimotor tasks, contributing to the growing body of evidence regarding WM’s involvement in cognitive functions. Our comprehensive analysis highlights the functional significance of WM ICNs and the need for continued exploration of WM’s contributions to cognitive processes.

### Clinical Implications: Unveiling Potential Biomarkers for Schizophrenia

The significant differences in functional connectivity observed between SZ patients and HC underscore the potential for developing more effective diagnostic tools and treatment strategies for SZ. Our findings, illustrated in the T-statistic matrix (Figure 11), reveal distinct alterations in both GM and WM connectivity patterns. The sFNC t-value matrix highlights a marked reduction in WM connectivity among SZ patients, particularly in the second WM module, with blue regions indicating significant negative t-values (SZ<HC). This reduction in connectivity was particularly pronounced in regions such as the insular, temporoparietal, and subcortical extended thalamus, suggesting disruptions in crucial white matter pathways that facilitate efficient communication between brain regions. Moreover, we observed notable inter-WM domain differences, with decreased connectivity between the frontal and subcortical basal ganglia, frontal and temporoparietal, frontal and insular, and frontal and sensorimotor regions. This widespread decline in cross-domain WM connectivity in SZ highlights the intricate nature of the disorder’s neurobiology. Conversely, the GM matrix exhibited a mixed pattern, with both red (indicating increased connectivity) and blue regions. Specific areas, such as the sensory-motor and subcortical-extended thalamus (SC-ET) regions, showed hyperconnectivity in SZ patients compared to HC. This heightened activity in certain GM regions may reflect compensatory mechanisms or altered functional processing, potentially serving as an adaptive response to the underlying disruptions in WM connectivity. The combined GM and WM sFNC matrix further elucidates these complex interactions, particularly in the first and third quadrants, where increased connectivity between WM frontal regions and GM areas related to higher cognition, insular, and temporal functions was observed in SZ patients. This intricate interplay between WM disruptions and GM hyperactivity suggests a broader dysfunction in brain connectivity, highlighting the potential for these altered connectivity patterns to serve as biomarkers for SZ diagnosis and treatment (Dini et al., 2024; Hassanzadeh et al., 2024). While previous studies have reported abnormalities in white matter microstructure in SZ patients using techniques like diffusion tensor imaging (DTI) (Chiapponi et al., 2013; S.-H. Lee et al., 2013; Szeszko et al., 2005), they primarily focused on global metrics or specific white matter tracts. In contrast, our network-level perspective provides novel insights into disrupted connectivity patterns within WM ICNs of SZ patients, aligning with the findings of Dini et al. (2024), who identified aberrant functional connectivity in resting-state networks among SZ patients. These insights point to the potential of utilizing altered connectivity patterns as biomarkers for more tailored diagnostic and therapeutic approaches, ultimately enhancing our understanding of the neurobiological basis of schizophrenia.

### Future Directions

The current findings open exciting avenues for future research. Exploring WM ICNs in various neurological and psychiatric disorders can provide valuable insights into their broader contributions to brain health and disease. Additionally, combining WM ICNs with other imaging modalities, such as diffusion MRI, holds immense promise for elucidating the intricate relationships between structure and function in the brain (Tang et al., 2017). Investigation of the potential for using WM ICNs to monitor treatment response or disease progression also holds promise for developing more targeted therapeutic interventions. Furthermore, expanding datasets to encompass diverse populations, including individuals of different ages, ethnicities, and socioeconomic backgrounds, is crucial for improving the generalizability and applicability of WM ICNs. Future research should aim to elucidate the mechanisms underlying WM’s involvement in task-related activities and explore its potential implications in neurodevelopmental and neurodegenerative disorders. Understanding the intricate interplay between WM and GM could provide new avenues for therapeutic interventions and improve our comprehension of neural dynamics in health and disease. This will ultimately lead to a more comprehensive model of brain connectivity that considers the complex interplay between WM and GM networks. Moreover, integrating WM ICNs with genetic and behavioral data could further elucidate the mechanisms underlying various brain disorders, leading to more effective treatments and interventions. In summary, this study highlights the novelty and significance of a WM ICN template. Our findings contribute to a deeper understanding of WM connectivity and its clinical relevance, particularly in relation to SZ. The distinct functional connectivity patterns observed in WM and GM ICNs underscore the importance of considering both in neuroimaging research. This work calls for ongoing research and collaboration to further unravel the complexities of brain function and dysfunction through the study of WM ICNs. The open-source availability of our template will support its integration into existing neuroimaging frameworks, promoting further exploration and discovery in the field.

## Conclusion

In summary, this study introduces a novel WM template with 97 ICNs and an approach for investigating WM functional connectivity, leveraging advanced techniques like scICA. By exploring both resting-state and task-based fMRI data, we provide a comprehensive framework for understanding the functional connectivity and spectral properties within WM as well as WM’s implications in SZ and HC. This work represents a significant advancement in neuroimaging, offering new tools and insights for future research into WM brain connectivity and its role in the overall functional architecture of the brain.

## Acknowledgments

Vaibhavi Itkyal, Armin Iraji, and Vince Calhoun contributed to the conception and design of the study. Data collection and pre-processing were performed by Vaibhavi Itkyal, Armin Iraji, Zening Fu and Jill Fries. Vaibhavi Itkyal is the first author and completed the data post-processing, experiments, visualization and wrote the first draft of the manuscript. All other authors made edits and suggestions to the manuscript. The project was completed under the supervision of lab director, Dr. Vince Calhoun. All authors commented on previous versions of the manuscript. All authors read and approved the final manuscript. This study was supported by NIH grant # R01MH123610, #5R01MH119251 and NSF grant #2112455.

## Supplementary figures

**Supplementary Figure 1:**
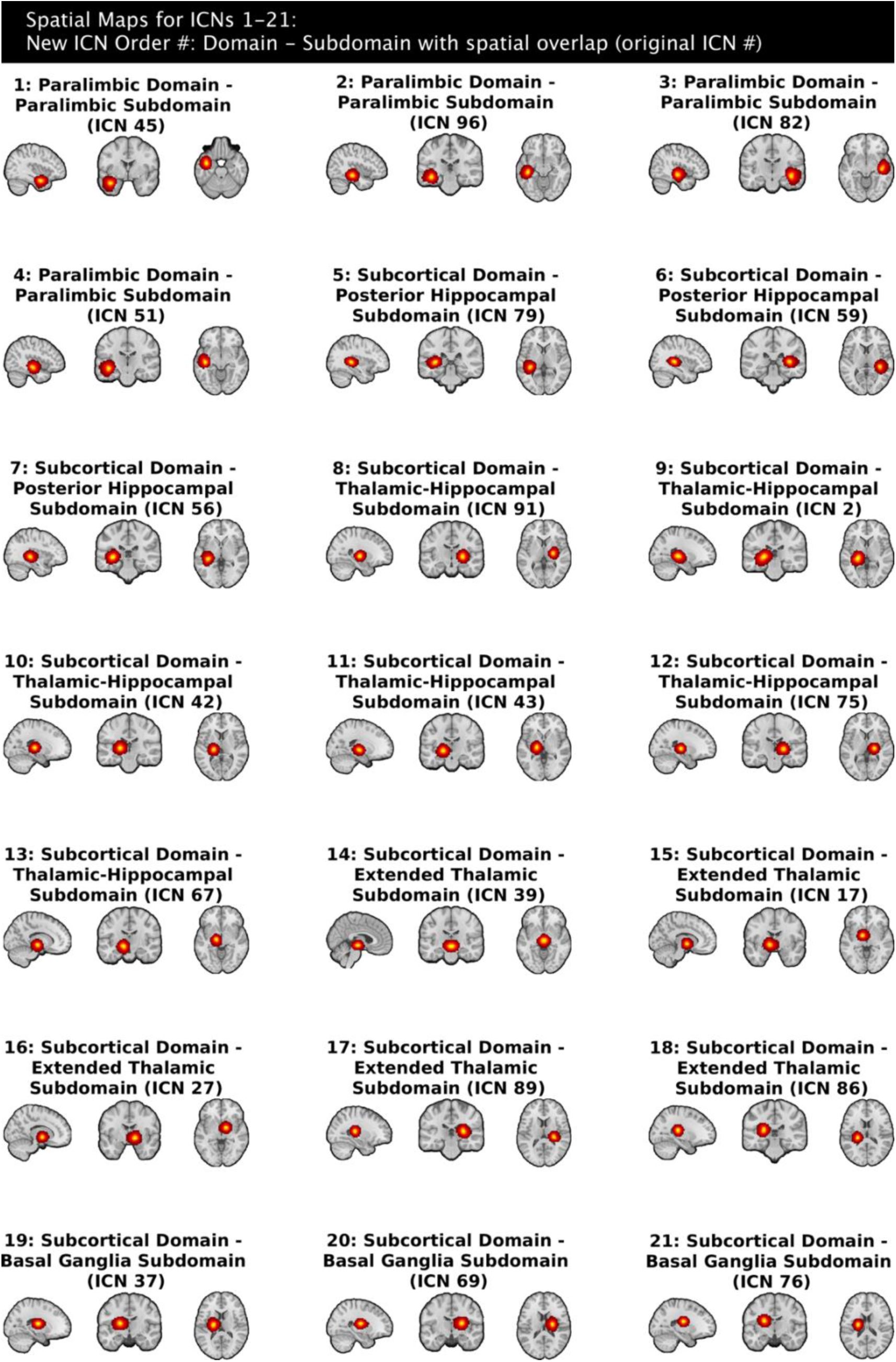
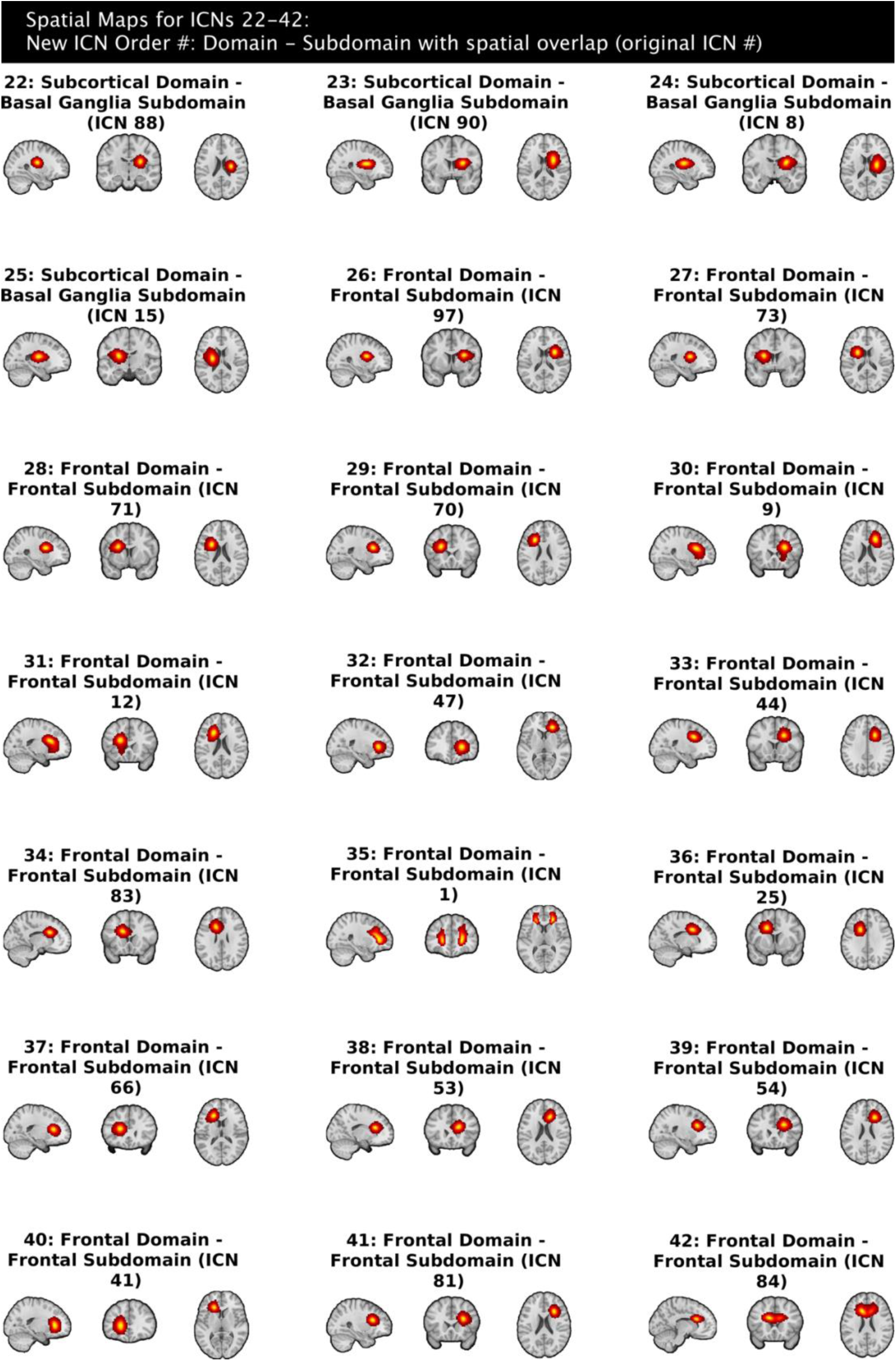
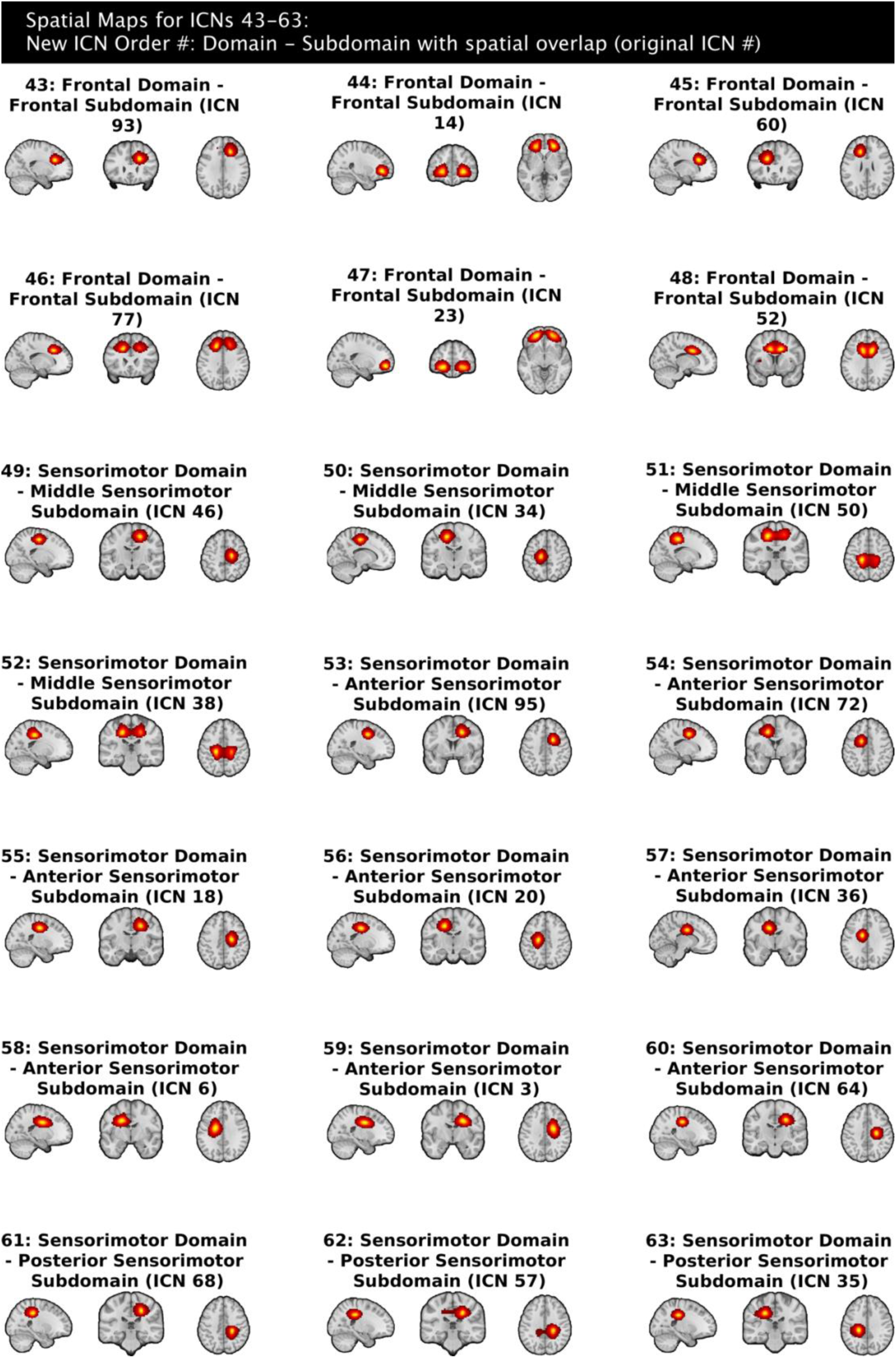
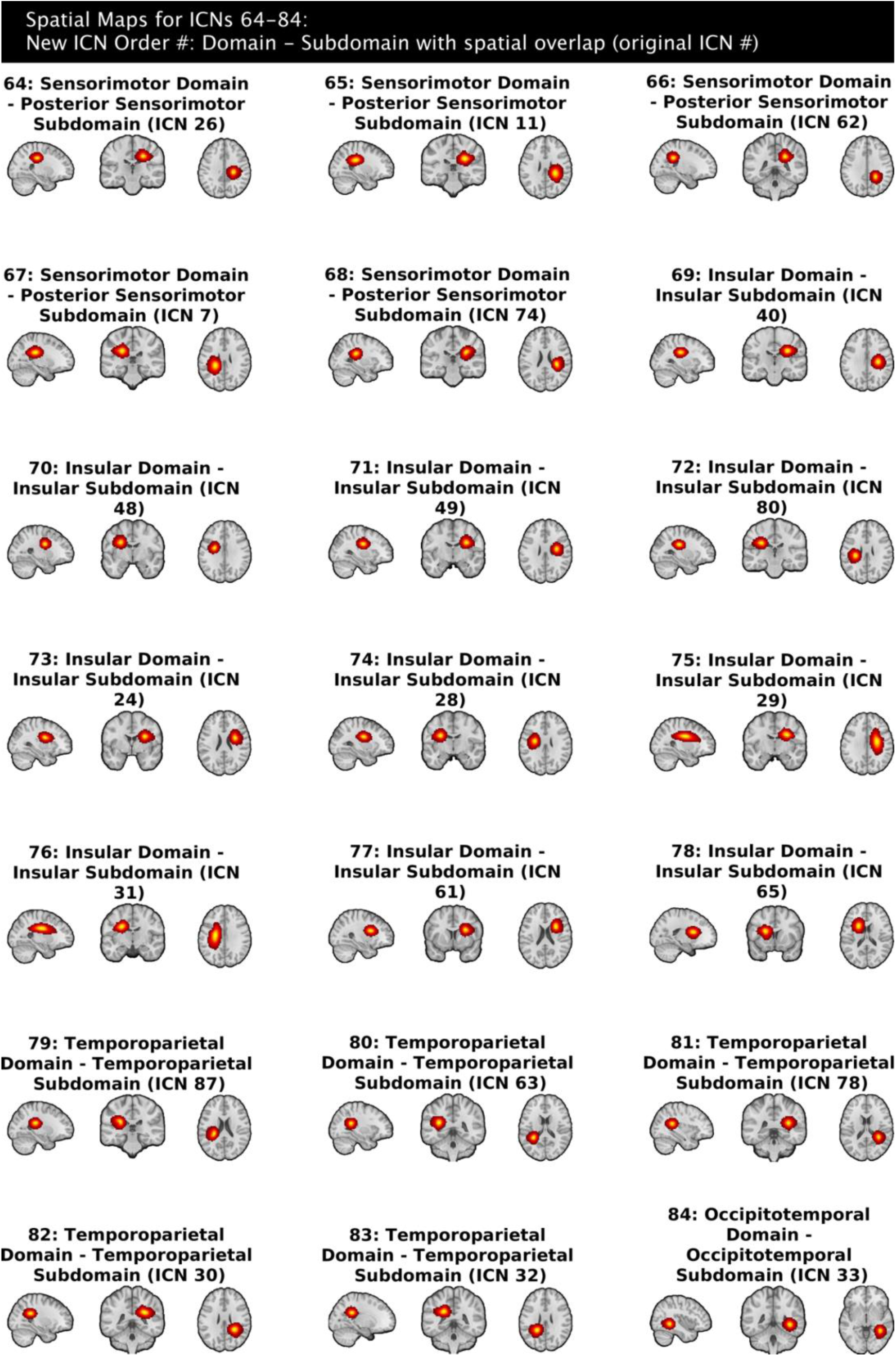
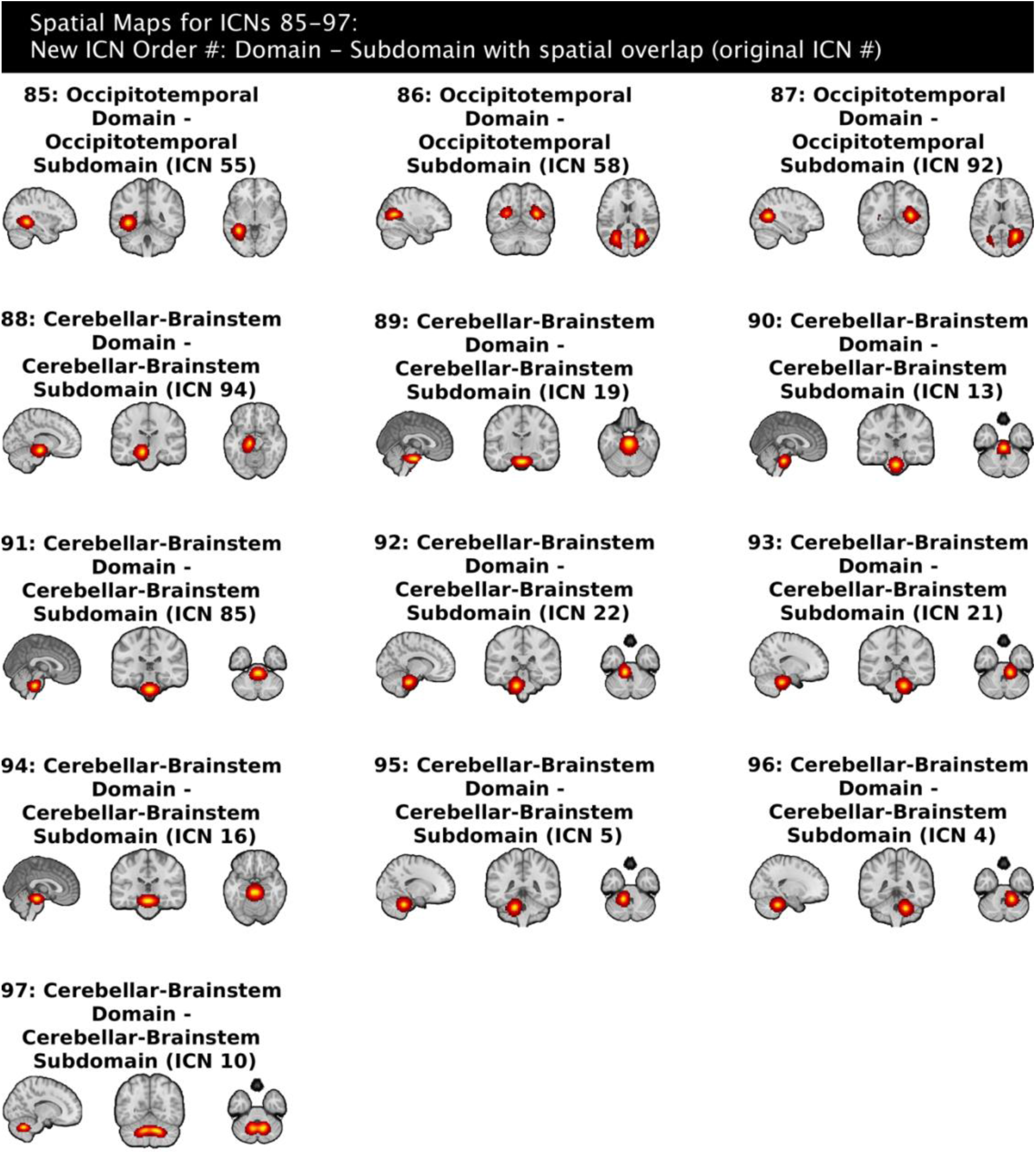
Individual WM ICN spatial maps.

